# Detecting local Zika virus transmission in the continental United States: a comparison of surveillance strategies

**DOI:** 10.1101/145102

**Authors:** Steven P Russell, Kyle R Ryff, Carolyn V Gould, Stacey W Martin, Michael A Johansson

**Author notes:** The findings and conclusions in this report are those of the authors and do not necessarily represent the official position of the Centers for Disease Control and Prevention.

## Abstract

**Introduction:** The 2015-2017 Zika virus (ZIKV) epidemic in the Americas has driven efforts to strengthen surveillance systems and to develop interventions, testing, and travel recommendations. In the continental U.S. and Hawaii, where limited transmission has been observed, detecting local transmission is a key public health objective. We assessed the effectiveness of three general surveillance strategies for this situation: testing all pregnant women twice during pregnancy, testing blood donations, and testing symptomatic people who seek medical care in an emergency department (ED).

**Methods:** We developed a simulation model for each surveillance strategy and simulated different transmission scenarios with varying population sizes and infection rates. We then calculated the probability of detecting transmission, the number of tests needed, and the number of false positive test results.

**Results:** The probability of detecting ZIKV transmission was highest for testing ED patients with Zika symptoms, followed by pregnant women and blood donors, in that order. The magnitude of the difference in probability of detection between strategies depended on the incidence of infection. Testing ED patients required fewer tests and resulted in fewer false positives than surveillance among pregnant women. The optimal strategy identified was to test ED patients with at least two Zika virus disease symptoms. This case definition resulted in a high probability of detection with relatively few tests and false positives.

**Discussion:** In the continental U.S. and Hawaii, where local ZIKV transmission is rare, optimizing the probability of detecting infections while minimizing resource usage is particularly important. Local surveillance strategies will be influenced by existing public health system infrastructure, but should also consider the effectiveness of different approaches. This analysis demonstrated differences across strategies and indicated that testing symptomatic ED patients is generally a more efficient strategy for detecting transmission than routine testing of pregnant women or blood donors.

## Introduction

In 2015 and 2016, Zika virus (ZIKV) spread through the Americas, with the first cases confirmed in early 2015 in Brazil [1] and 48 countries and territories reporting confirmed locally acquired infections by the end of 2016 [2]. Prior to its emergence in the Americas, ZIKV was a relatively obscure arboviral disease, with the first documented outbreak occurring on the island of Yap in 2007 [3]. While ZIKV infection appeared to be relatively benign in the Yap outbreak, the study of subsequent outbreaks in French Polynesia and the Americas provided evidence that ZIKV infection was associated with adverse outcomes, including Guillain-Barré syndrome [4–6] and congenital birth defects [7]. These severe manifestations have driven efforts to identify areas where transmission is ongoing to target interventions, testing, and travel recommendations.

Many areas of the Americas were also impacted by the emergence of chikungunya virus in 2013-2014 and have endemic dengue virus transmission [8], two arboviruses transmitted by the same *Aedes* mosquito vectors. These areas were at risk of ZIKV transmission and most experienced large Zika outbreaks. In the continental United States, however, Zika, chikungunya, and dengue viruses have each been repeatedly introduced into areas inhabited by *Ae. aegypti* and *Ae. albopictus*, yet transmission has only been sporadically identified [9–13]. The climate of the southern U.S. is suitable for Zika virus transmission but living conditions (e.g., air conditioning) restrict human-mosquito interaction, likely limiting transmission [14].

Although the risk of widespread mosquito-borne transmission in the continental U.S. and Hawaii is low, locally acquired cases have been reported in Florida and Texas [12, 13]. In August 2016, the Food and Drug Administration (FDA) issued revised recommendations for testing all donated blood products in the United States [15]. These recommendations were implemented to protect the blood supply but could also serve as a surveillance strategy to detect asymptomatic or pre-symptomatic Zika virus infections and monitor transmission as has been done in Puerto Rico [16]. However, this is not the only possible strategy for detecting local ZIKV infections in areas at risk for mosquito-borne transmission [17]. Locally implemented surveillance systems should consider several factors: the sensitivity (probability of detecting transmission should it occur), the specificity (avoidance of false-positive test results requiring follow-up), and the resources needed for conducting surveillance (e.g. laboratory testing capacity).

We assessed the likely effectiveness of three general surveillance strategies that could be used to detect ZIKV transmission in areas of the continental U.S. and Hawaii at risk of local mosquito-borne transmission. The first strategy is to test all pregnant women twice during pregnancy (1^st^ and 2^nd^ trimester), as was previously recommended in areas with known ongoing risk [18]. This would align surveillance with the population of greatest concern, but it limits testing to that group and it is possible that transmission could be detected earlier in other populations. The second strategy is to use the results of blood donation screening, which is already being conducted as recommended by the FDA. Blood donation is only permitted for healthy individuals, so this strategy would only detect asymptomatic or pre-symptomatic infections. The population tested is also limited to the population seeking to donate; the U.S. averages about 4.3-4.7 blood donations per 100 people per year [19, 20]. The third strategy is to test symptomatic people who seek medical care. In Yap and French Polynesia, serosurvey results suggest that only a small proportion of infected individuals were symptomatic and sought care [3, 21, 22] and symptoms were possibly non-specific. Nonetheless, there may be a higher probability of detecting infections among people seeking care, which we limit to emergency department (ED) visits for the purposes of the current study.

## Methods

We first identified available data for critical components of each of the three surveillance strategies, which are described below and summarized in Table 1. For the purposes of this assessment, pregnant women were assumed to be tested solely with a serological assay to detect anti-ZIKV IgM, which is currently included in the algorithm for asymptomatic pregnant women with ongoing risk for exposure to Zika virus as part of routine obstetric care [18]. Data on Zika IgM ELISA test sensitivity and specificity are limited, but based on studies assessing IgM ELISA test performance for dengue viruses [23–26], we assumed the test would have a sensitivity of 80-99%. In the context of the continental U.S., where dengue is not endemic and the potential cross-reactivity of related flaviviruses is of limited concern, we assumed a specificity of 80-95%. We estimated that IgM antibodies would be detectable in an infected person for a period of 2-4 months [27–29]. Live birth rates across U.S. states vary from 9 to 17 per 1,000 persons per year [30]. We assumed that testing would occur only in pregnant women (excluding pregnancies that do not reach full term) and that women would be tested twice during pregnancy, in the first and second trimesters.

**Table 1.** Parameter assumptions for the three surveillance strategies

| Parameter | Estimate | Source |
| --- | --- | --- |
| **Pregnant women** |  |  |
| Sensitivity of IgM MAC ELISA<sup>i</sup> | 80-99% | [23-26] |
| Specificity of IgM MAC ELISA<sup>i</sup> | 80-95% | [23-26] |
| Time that IgM antibodies are detectable | 2-4 months | [27-29] |
| Birth rate | 9-17 per 1,000 persons per year | [30] |
| **Blood bank donors** |  |  |
| Sensitivity of BDS-NAT<sup>ii</sup> test | 98-100% | [34] |
| Specificity of BDS-NAT<sup>ii</sup> test | 99.99-100% | [31, 32] |
| Time that virus is detectable | 11-17 days | [28] |
| Blood donation rate | 43-47 per 1,000 persons per year | [19, 20] |
| Percentage of ZIKV infections that are asymptomatic | 60-80% | [3, 21, 22] |
| **ED patients** |  |  |
| Sensitivity of RT-PCR<sup>iii</sup> test | 80-95% | [28] |
| Specificity of RT-PCR<sup>iii</sup> test | 99-100% | [35] |
| Percentage of people who visit an ED in a given week | 0.7%-1% | [37] |
| Of those, percentage who had: |  |  |
| Fever | 3.7-6.7% | [38] |
| Headache | 2.7-3.6% | [38] |
| Rash | 4.3-6.1% | [38] |
| Arthralgia | 0.8-1.4% | [38] |
| Conjunctivitis | 0.1-0.3% | [38] |
| Percentage of ZIKV infected people who have symptoms | 20-40% | [3, 21, 22] |
| Of those, percentage who sought care | 10-50% | [3, 21, 22] |
| Of those, percentage who sought care at an ED<sup>iv</sup> | 5-50% | [39] |
| Of those, percentage who had: |  |  |
| Fever | 51-53% | [36] |
| Headache | 65-67% | [36] |
| Rash | 86-87% | [36] |
| Arthralgia | 60-61% | [36] |
| Conjunctivitis | 16-17% | [36] |

<sup>i</sup>Immunoglobulin M antibody capture enzyme-linked immunosorbent assay
<sup>ii</sup>Nucleic acid test
<sup>iii</sup>Reverse transcription polymerase chain reaction
<sup>iv</sup>Emergency Department

In the context of surveillance, we assumed that blood donations would be tested with highly specific blood donor screening nucleic acid tests (BDS-NAT) such as the cobas [31] and Procleix [32] assays. We assumed that these assays could detect ZIKV RNA for 11-17 days, the median time to ZIKV RNA clearance [28], and would have a specificity of greater than 99.99, using a range of 99.99-100% [31, 32]. Clinical sensitivity of the these assays has not been characterized, but analytical sensitivity is higher than typical RT-PCR tests [33] and we assumed it would be in the range of 98-100% based on clinical sensitivity estimates for the West Nile virus individual NAT assay [34]. A recent study estimated that approximately 13,639,000-14,835,000 whole blood and apheresis red blood cell units were collected in the U.S. in 2013 [19]. With a population of 316,128,839 U.S. residents [20], the estimated rate of donation was 43-47 donations per 1,000 persons per year. We assumed that 60-80% of individuals infected with ZIKV would be asymptomatic and therefore eligible for blood donation [3, 21, 22].

For the purposes of this study, we assumed that symptomatic ED patients would be tested exclusively by serum RT-PCR, with no IgM testing. Since ED patients typically present while symptomatic and viral RNA is estimated to be detectable for a median of 11-17 days [28], we assumed that all ZIKV infected ED patients would have detectable RNA. We assumed that the RT-PCR test has a sensitivity of 80-95%, based on the confidence interval for 118 out of 134 patients who tested positive by RT-PCR within 7 days of symptom onset in a recent study [28]. We assumed that the specificity of the RT-PCR test would be greater than 99% and used a range of 99-100% [35]. We focused on patients with five symptoms associated with ZIKV infection: fever, headache, rash, arthralgia and conjunctivitis [36]. First, we estimated that 0.7%-1% of the population visits an ED in a given week [37]. Then, we used data extracted from the National Syndromic Surveillance Program’s BioSense Platform [38] on aggregated chief complaints for 1-2 million ED visits on a weekly basis over the period from the week starting May 29, 2015 to the week starting May 28, 2016. Only national-level aggregate data from U.S. states were used. For each symptom and combination of symptoms, we found the range of weekly frequencies at which that symptom or combination was reported. For example, across all the weeks, a minimum of 3.6% and a maximum of 6.7% of ED patient records included fever in the chief complaint description. Ranges for each symptom and combination of symptoms are provided in Supplemental Table 1. We then estimated the probability of ZIKV-infected individuals seeking care in the EDs. We assumed that 20-40% of ZIKV infections will exhibit symptoms [3, 21, 22], 10-50% of people with symptomatic infections will seek care [3, 21, 22], and 5-50% of care-seeking individuals will visit an ED [39]. Among those patients, we assumed that symptoms would be distributed with the same proportions found among symptomatic ZIKV-infected people seeking care in Puerto Rico [36] (Supplemental Table 1).

To account for uncertainty and variability in each parameter, we used the range of likely values to create uniform sampling distributions and drew 10,000 samples of each parameter from its respective distributions under transmission scenarios and population sizes using the following formula to calculate *P*_*detect*_, the weekly probability of detecting at least one ZIKV infection:

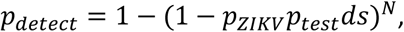

Where *P*_*ZIKV*_ is the weekly probability of being infected with ZIKV, *P*_*test*_ is the weekly probability of being tested, *d* is the duration (in weeks) of detectible RNA or antibodies, *s* is the sensitivity of the assay, and *N* is the population size. *P*_*zikv*_ and *N* were fixed for individual simulations, with *P_zikv_* ranging from 1 infection per 100,000 per week to 1 infection per 1,000 per week and *N* fixed at 10,000, 100,000, or 1,000,000. The assay specific variables, *d* and *s*, were simulated from uniform distributions over the range specified in Table 1 with the exception that *d* was assumed to be 1 for ED patients, as they would only be tested when presenting with symptoms. For pregnant women, *P*_*test*_ is twice the number of new pregnancies per week (to account for two tests per pregnancy) divided by the total population size. For blood bank donors, *P*_*test*_ is the product of the weekly probability of blood donation per person and the probability of infection being asymptomatic. For ED patients, *P*_*test*_ is the product of the proportion of ZIKV infected people who have any symptoms, the proportion of those who seek care at an ED, and the proportion of those who have each specific symptom or set of symptoms.

For each strategy, the expected number of tests required, *T*, is a binomial sample from the population, *N*, with probability, *p*_*test*_. The expected number of false positive tests is a binomial sample with probability 1 – *specificity* from the total number of tests, *T*. The proportion of total infections detected is *p*_*test*_*ds*. We summarized the results using the 25^th^ and 75^th^ percentiles to identify 50% uncertainty intervals, and the 2.5^th^ and 97.5^th^ percentiles to identify 95% uncertainty intervals.

## Results

We analyzed three general surveillance strategies: testing asymptomatic pregnant women, testing blood donors, and testing ED patients with either rash (the most common Zika symptom) or rash and headache (the most common combination of Zika symptoms, Supplemental Table 1). Regardless of population size and infection rate, the probability of detecting at least one ZIKV infection was highest for testing ED patients with rash, followed by ED patients with both rash and headache, pregnant women, and blood donors, in that order (Figure 1). In the smallest population considered (10,000 people), the weekly probability of detection did not reach 25% for any system, even with weekly infection rates as high as 1 ZIKV infection per 1,000 people. In larger populations, the weekly probability of detection increased as the total number of infections increased. In a population of 100,000 people with an incidence rate of 1 ZIKV infection per 1,000 people (or equivalently, in a population of 1 million people with an incidence rate of 1 ZIKV infection per 10,000 people), testing ED patients with rash resulted in a weekly probability of detection of 79% (50% Uncertainty Interval (UI): 59%, 93%). In the same population, the probability of detection was 66% (50% UI: 46%, 83%) when testing ED patients with both rash and headache, 40% (50% UI: 35%, 47%) when testing pregnant women, and 11% (50% UI: 10%, 12%) when testing blood donors. In a population of 1 million with an incidence of 1 ZIKV infection per 1,000 people, testing pregnant women or ED patients in either group resulted in probabilities of detection higher than 99%, while testing blood donors resulted in a probability of detection of 70% (50% UI: 65%, 74%). Supplemental Figure 1 shows the 95% uncertainty intervals for these estimates.

**Figure 1.**
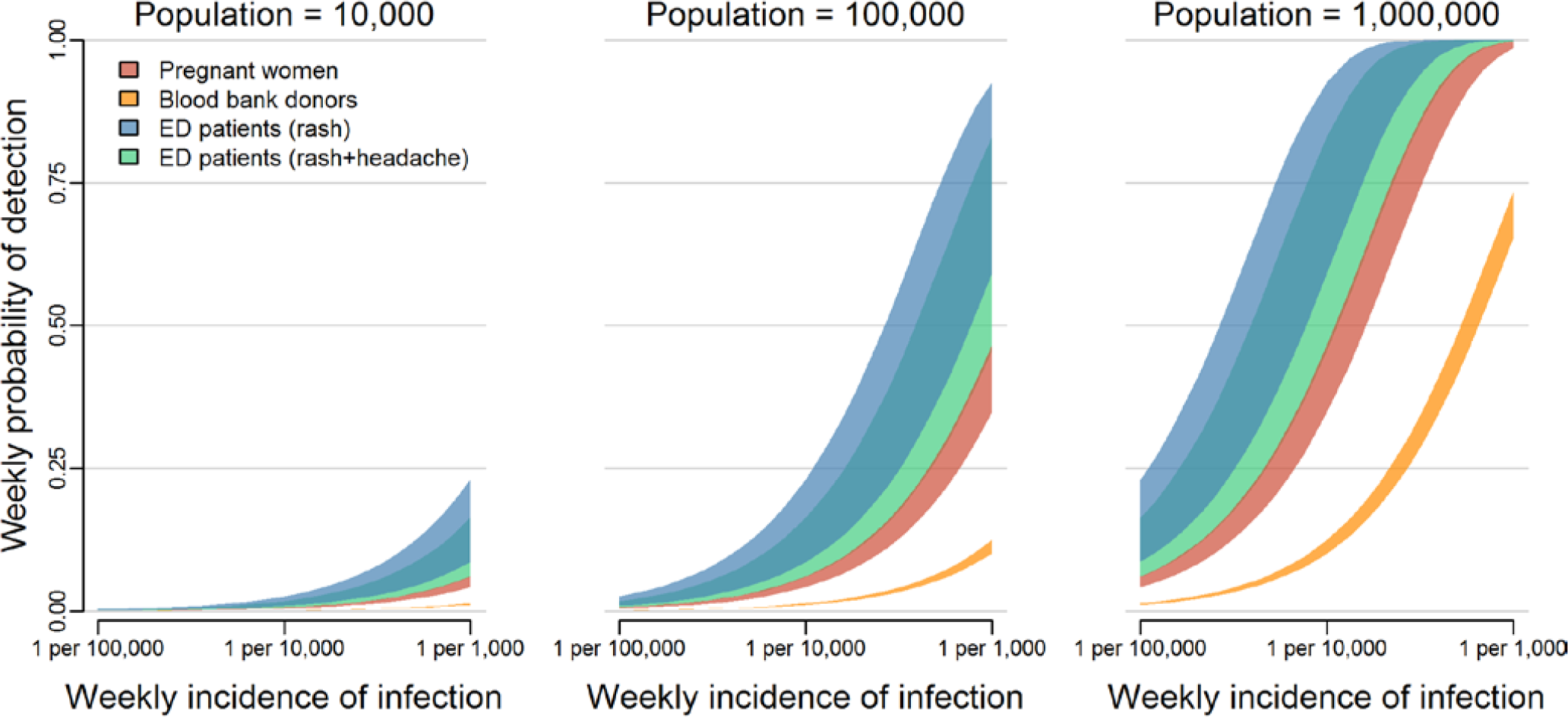
The probability of detecting local transmission for different surveillance strategies. The overlapping bands represent 50% uncertainty intervals for the weekly probability of detecting transmission by testing asymptomatic pregnant women (red), blood donors (orange), or patients in emergency departments exhibiting rash (blue) or rash and headache (green). These probabilities are shown for three population sizes over a range of possible ZIKV incidences. Supplemental Figure 1 shows the 95% uncertainty intervals.

The expected number of tests also varied between systems (Figure 2A). In a population of 100,000, surveillance among blood donors required an estimated 87 (95% UI: 83, 90) BDS-NAT tests per week regardless of ZIKV incidence; these tests are already performed as routine screening of the blood supply. Routine testing of all pregnant women would require 50 (95% UI: 35, 65) IgM ELISA tests per week. Under the high incidence scenario (1 infection per 1,000 people per week), testing ED patients with rash would require 46 (95% UI: 35, 59) RT-PCR tests per week and testing ED patients with rash and headache would require only 2.5 (95% UI: 1.4, 5.2) RT-PCR tests per week. For smaller and larger populations, the number of tests scaled directly with population sizes. For example, in a population of 1 million people, 870 (95% UI: 830, 900) BDS-NAT tests would be expected for blood donor surveillance (not shown). The number of tests performed does not correlate exactly with the probability of detection. For example, while fewer tests are required for surveillance among pregnant women compared to blood donors, the probability of detection was higher because of the longer detection window of IgM antibodies compared to ZIKV RNA.

**Figure 2.**
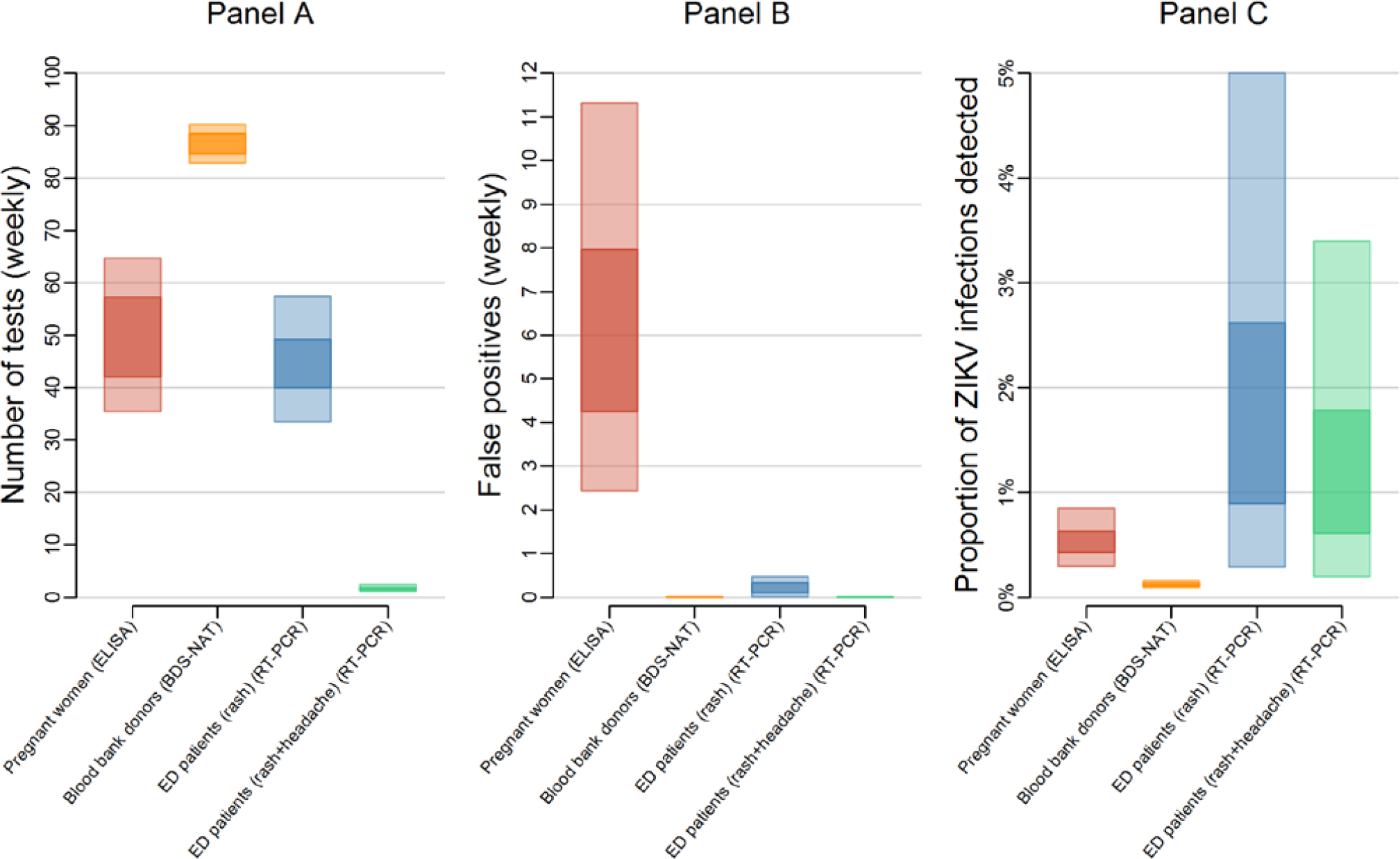
Expected tests, false positives, and proportion of ZIKV infections detected for different surveillance strategies. Panel A shows the 50% uncertainty interval (UI, dark) and 95% UI (light) for the expected number of tests needed per week to test pregnant women (red), blood bank donations (orange), patients in emergency departments exhibiting rash (blue), and patients in emergency departments exhibiting rash and headache (green) in a population of 100,000 people. Panel B shows the estimated weekly number of false positives in a population of 100,000 people. The number of tests and false positives (Panels A and B) for pregnant women and blood donor testing do not change relative to ZIKV incidence. Estimates for ED patients reflect the relatively high transmission scenario of 1 infection per 1,000 people per week. Panel C shows the proportion of true ZIKV infections that would be detected if ZIKV transmission was occurring.

With little or no local transmission, tests are performed mostly on people who are not infected and some of those tests result in false positives, with the number of false positives being dependent on the specificity of the assay. The assay with the lowest specificity was the IgM ELISA, which was considered for surveillance among pregnant women. This resulted in the high median number of false positive test results — 6.0 (95% UI: 2.4, 11.0) per week in a population of 100,000 (Figure 2B). In contrast, both the BDS-NAT and RT-PCR assays are highly specific and have reduced rates of false positives compared to the IgM ELISA assay. Testing ED patients with rash would result in an estimated 0.22 (95% UI: 0.01, 0.47) false positive tests per week, while testing ED patients with both rash and headache would result in an estimated 0.006 (95% UI: 0.0003, 0.01) false positives per week, and testing blood donors with the BDS-NAT assay would result in an expected 0.004 (95% UI: 0.0002, 0.008) false positive tests per week. In strategies that use a limited number of highly specific tests, a positive result can accurately predict a true infection. Even at low infection rates, a single test administered on an ED patient with rash and headache had a high positive predictive value (Figure 3, Supplemental Figure 2).

**Figure 3.**
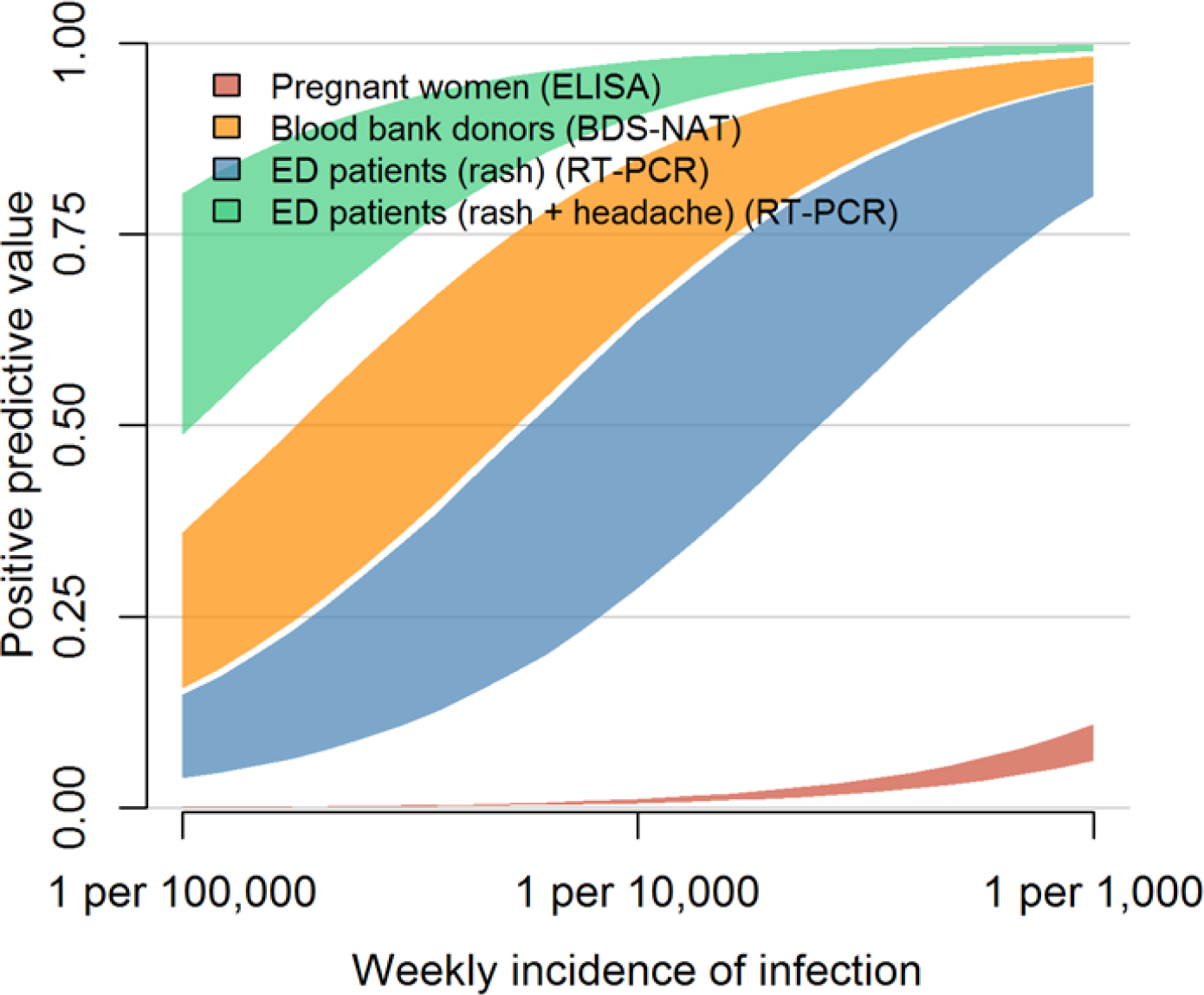
Positive predictive value for different surveillance strategies. This figure describes the positive predictive value (PPV) of a single positive test result under different surveillance strategies: testing asymptomatic pregnant women (red), blood donors (orange), ED patients exhibiting rash (blue) or ED patients exhibiting rash and headache (green). The bands represent 50% uncertainty intervals for the PPV of a positive test in each surveillance system over a range of possible ZIKV incidences; Supplemental Figure 2 shows 95% uncertainty intervals.

The expected proportion of infections detected in any strategy was low. The highest proportion of detection occurred when testing all ED patients with rash, in which 1.6% of all ZIKV infections (95% UI: 0.29, 5.0) would be detected (Figure 2C). Testing ED patients with rash and headache was second highest, with 1.1% of infections detected (95% UI: 0.2, 3.4), followed by pregnant women, 0.5% of infections (95% UI: 0.3%, 0.9%), and blood donors, 0.1% of infections (95% UI: 0.1%, 0.2%). ED patient testing resulted in a higher probability of detection, fewer tests, and fewer false positives, so we investigated alternative case definitions for surveillance. The number of RT-PCR tests required (Figure 4A) depends on the number of ED patients who meet each case definition. Among Zika-like symptoms in ED patients that we analyzed, fever, rash, and headache were most commonly reported; 3.7-6.7% reported fever, 4.3-6.1% reported rash, and 2.7-3.6% reported headache (Supplemental Table 1). Combinations of these symptoms were rare, 0.9-1.6% had two or more of these five symptoms and 0.3-0.7% had three or more of these five symptoms. Additionally, 0.6-0.9% has rash plus at least one of the other four symptoms.

**Figure 4.**
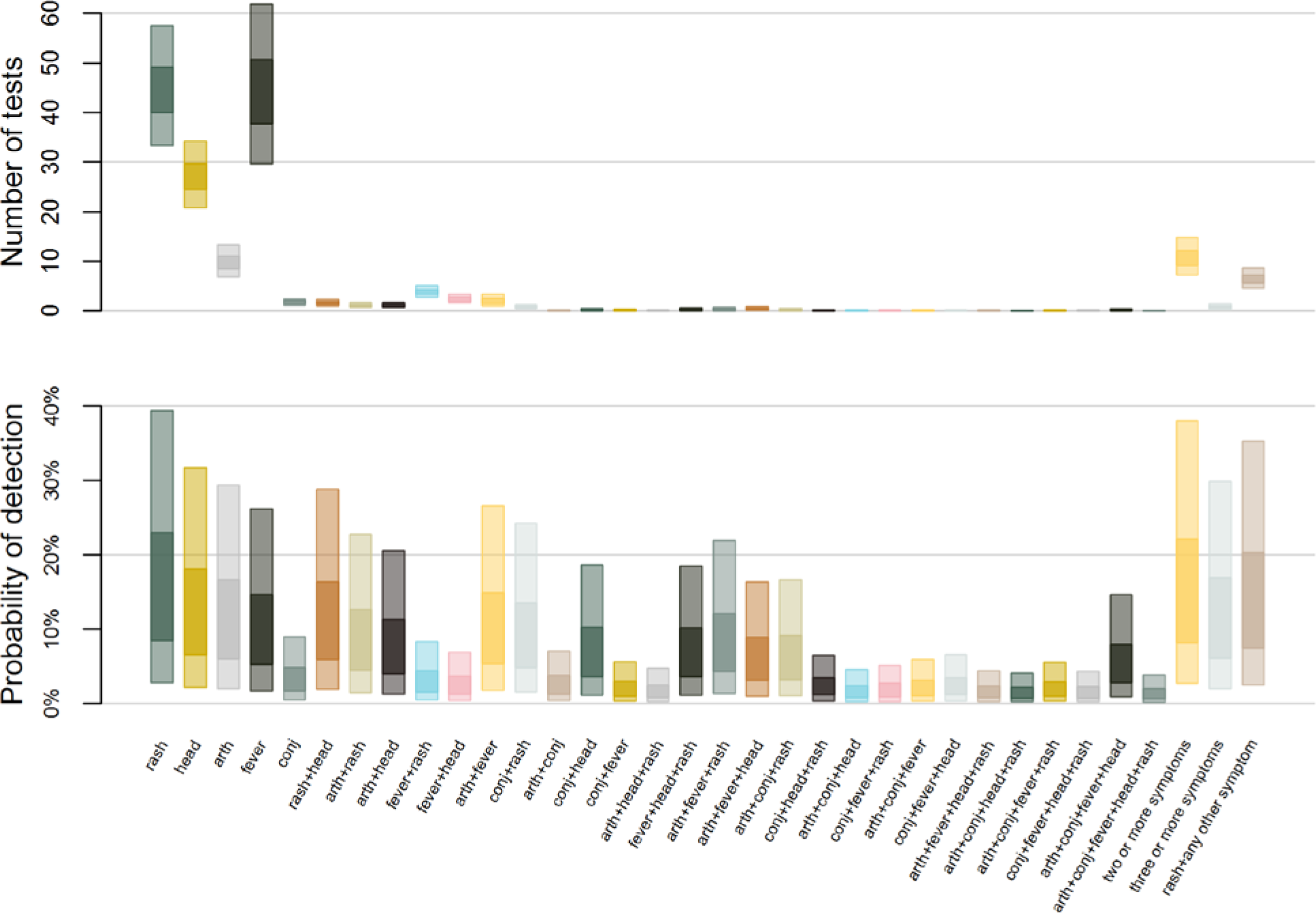
Estimated number of tests and corresponding probability of detection by case definition. The top plot shows the estimated weekly number of tests required to conduct surveillance using different case definitions in a population of 100,000 people with ZIKV incidence of 1 infection per 1,000 people per week. Case definitions include rash, headache (head), arthralgia (arth), fever, conjunctivitis (conj), and combinations of those five symptoms or rash plus at least one more. The bottom plot shows the corresponding probability of detecting local transmission using each case definition. The inner box represents 50% uncertainty while the outer box represents 95% uncertainty.

The surveillance case definition also determines the likelihood of testing and therefore the probability of detection (Figure 4B). Among ZIKV infected individuals, rash was the most common symptom (86-87%), followed by headache (65-67%), arthralgia (60-61%), fever (51-53%), and conjunctivitis (16-17%) (Supplemental Table 1). Multi-symptom combinations were also common; 82-83% of infected individuals had two or more of these five symptoms and 61-62% had three or more of these five symptoms. Additionally, 75-76% had rash plus at least one of the other four symptoms. Accordingly, the probability of detection was highest when testing for ED patients with rash, followed by testing patients with two or more symptoms, rash and one other symptom, and so on.

As shown in Figures 1 and 2, testing of symptomatic ED patients with rash outperformed testing of pregnant women with a higher probability of detection and fewer false positive test results. The alternative cases definitions of (a) at least two symptoms, (b) at least three symptoms, and (c) rash plus at least one additional symptom, were almost as effective at detecting transmission as testing anyone with rash (Figure 5A, Table 2). In a population of 100,000 people with a weekly incidence rate of 1 ZIKV infection per 10,000 people, the median estimated probability of detection for the multi-symptom case definitions was 11-14% compared to 15% for all patients with rash. Meanwhile, the multi-symptom case definitions required far fewer tests; an average of 0.3-0.9 (95% UI) tests per week for patients with three or more symptoms, 4.5-8.2 (95% UI) tests per week for patients with rash plus another symptom, and 7-15 (95% UI) tests per week for patients with two or more symptoms (Figure 5B).

**Figure 5.**
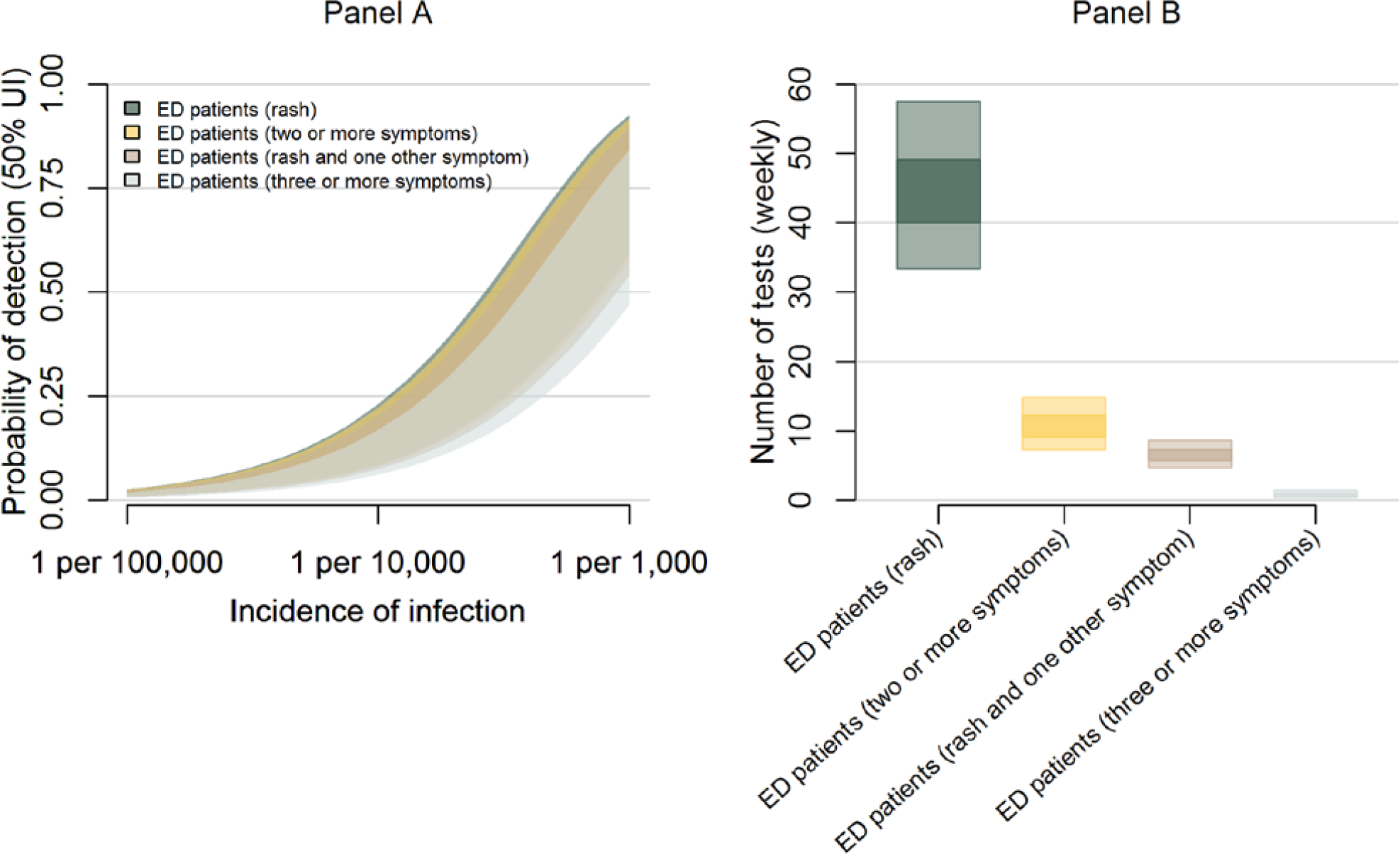
The probability of detection and testing burden for ED surveillance strategies. Panel A shows the weekly probability of detecting transmission with overlapping 50% uncertainty intervals for testing ED patients with rash (dark green), at least two symptoms (yellow), rash plus at least one additional symptom (brown), and at least three symptoms (grey). The probabilities are shown in a population of 100,000 over a range of possible ZIKV incidences. Panel B shows the estimated number of tests per week for each of these strategies with ZIKV incidence of 1 infection per 1,000 people per week.

**Table 2.** Estimated strategy performance (ZIKV incidence of 1 per 10,000 per week, population of 100,000)

| **Surveillance strategy** | **Weekly probability of detection (95% UI<sup>i</sup>)** | **Number of tests (95% UI<sup>i</sup>)** |
| --- | --- | --- |
| Pregnant women | 5% (2.9%, 8.2%) | 50 (35, 65) |
| Blood donors | 1.2% (0.9%, 1.5%) | 87 (83, 90) |
| ED<sup>ii</sup> patients (rash) | 15% (2.9%, 39%) | 44 (33, 57) |
| ED<sup>ii</sup> patients (rash + headache) | 10% (2%, 29%) | 1.4 (1.0, 1.9) |
| ED<sup>ii</sup> patients (two or more symptoms) | 14% (3%, 38%) | 10 (7.1, 14) |
| ED<sup>ii</sup> patients (three or more symptoms) | 11% (2%, 30%) | 0.57 (0.30, 0.92) |
| ED<sup>ii</sup> patients (rash + one other symptom) | 13% (3%, 35%) | 6.2 (4.5, 8.2) |

## Discussion

In areas of the US and Hawaii where there is potential risk of ZIKV transmission, detecting transmission should it occur is a key public health objective [17]. This poses a challenge for surveillance systems: to balance the probability of detecting ZIKV transmission with the resources needed to carry out surveillance when infections may be rare. An efficient surveillance system should have high sensitivity (a relatively high probability of detecting infections) and high specificity (a low probability of a positive test result in an individual without infection).

Comparing three general surveillance systems, we found that none of them was likely to detect even 5% of all infections that occur, reflecting the difficulty in detecting ZIKV transmission because a high proportion of infections are mild or asymptomatic. However, the most important feature of any surveillance system is not the ability to detect all cases, but the ability to detect at least one locally transmitted case and thus prompt public health authorities to conduct enhanced surveillance and recommend appropriate prevention measures in that location. Among these strategies, the system for testing individuals presenting in EDs had a higher probability of detecting at least one ZIKV infection than systems designed to detect infection in pregnant women or blood donors.

The numbers of tests required and false positive results were also lowest for ED patients with testing limited to RT-PCR. The number of tests was highest for blood donors, though this does not represent an added testing burden as blood donation screening is already implemented to protect the blood supply [15]. The number of false positive tests was highest among pregnant women because of the relatively low specificity of the IgM ELISA. In any system, the number of tests and false positive results will depend on the diagnostic testing algorithm that is used. For example, both IgM ELISA and RT-PCR testing could be implemented to increase sensitivity. For ED patients, the addition of IgM ELISA testing would increase the number of tests and false positives. The testing algorithm for pregnant women considered here also only represents one possible algorithm. Challenges with cross-reactivity and low specificity have prompted CDC to recommend a combination of diagnostic tests for pregnant women with exposure risk [40]. Complications of third trimester ZIKV infection have prompted recommendations for third trimester testing for those women [41]. If these changes were implemented in a surveillance system to detect transmission, the probability of detection, the number of tests, and the number of false positives would increase.

We found that the most common Zika symptom, rash, was also common among ED patients, being reported in 4-6% of ED visit chief complaints. However, far fewer ED patients met narrower case definitions of (i) two or more Zika symptoms (rash, headache, arthralgia, fever, or conjunctivitis), (ii) three or more Zika symptoms, or (iii) rash plus at least one other symptom. With these multiple symptom case definitions, we found that the probability of detection was almost equivalent to using rash alone, but the number of tests required and false positives were reduced substantially. This strategy thus represents a good combination of high probability of detection and low resource use.

This analysis only considered three general options for surveillance. It was intended not as an exhaustive review of all possibilities, but as a comparison of broadly applicable strategies with data that were available. The optimal strategy identified was to detect symptomatic infections with at least two Zika symptoms among patients presenting to the ED. However, this exact strategy may not be feasible nor optimal for any given location. For example, if RT-PCR testing is unavailable it would not be viable. If people do not tend to seek care in EDs, there may be alternative ways to try to capture cases. A jurisdiction may also choose to combine multiple approaches as part of a surveillance strategy. The analysis also did not consider the costs of assays or other resources (e.g. personnel, transportation, or laboratory supplies) needed to perform surveillance; costs could vary between systems and jurisdictions and therefore would be important to consider as part of any implementation plan. Furthermore, in areas where transmission has been detected and for travelers to areas of risk these strategies do not replace existing guidance for testing symptomatic persons or asymptomatic pregnant women [18].

The results presented here are general, but highlight key factors to consider and demonstrate how different approaches can be compared. Specifically, a surveillance system for local transmission of Zika virus in the continental U.S. and Hawaii is targeting a pathogen that is likely rare, such that extensive testing may be needed to detect transmission should it occur. In this situation, optimizing the probability of detecting infections while minimizing resource usage is particularly important. Directly comparing approaches through simulation, as done here, can highlight important tradeoffs and can inform a thorough analysis of potential surveillance strategies.

---

**Supplemental Figure 1.**
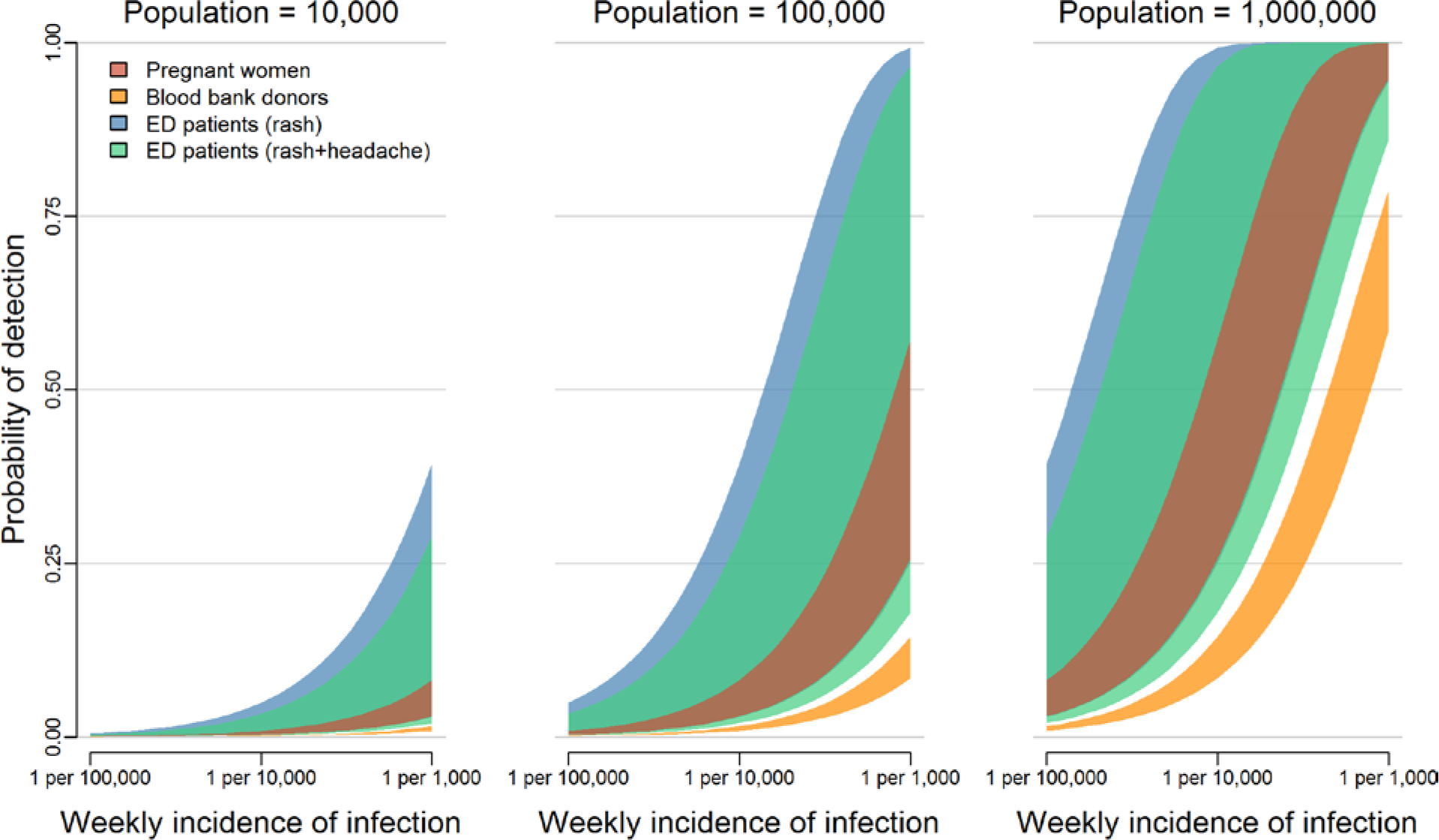
The probability of detecting local transmission for different surveillance strategies. The overlapping bands represent 95% uncertainty intervals for the weekly probability of detecting transmission by testing pregnant women (red), blood donors (orange), or patients in emergency departments exhibiting rash (blue) or rash and headache (green). These probabilities are shown for three population sizes over a range of possible incidences.

**Supplemental Figure 2.**
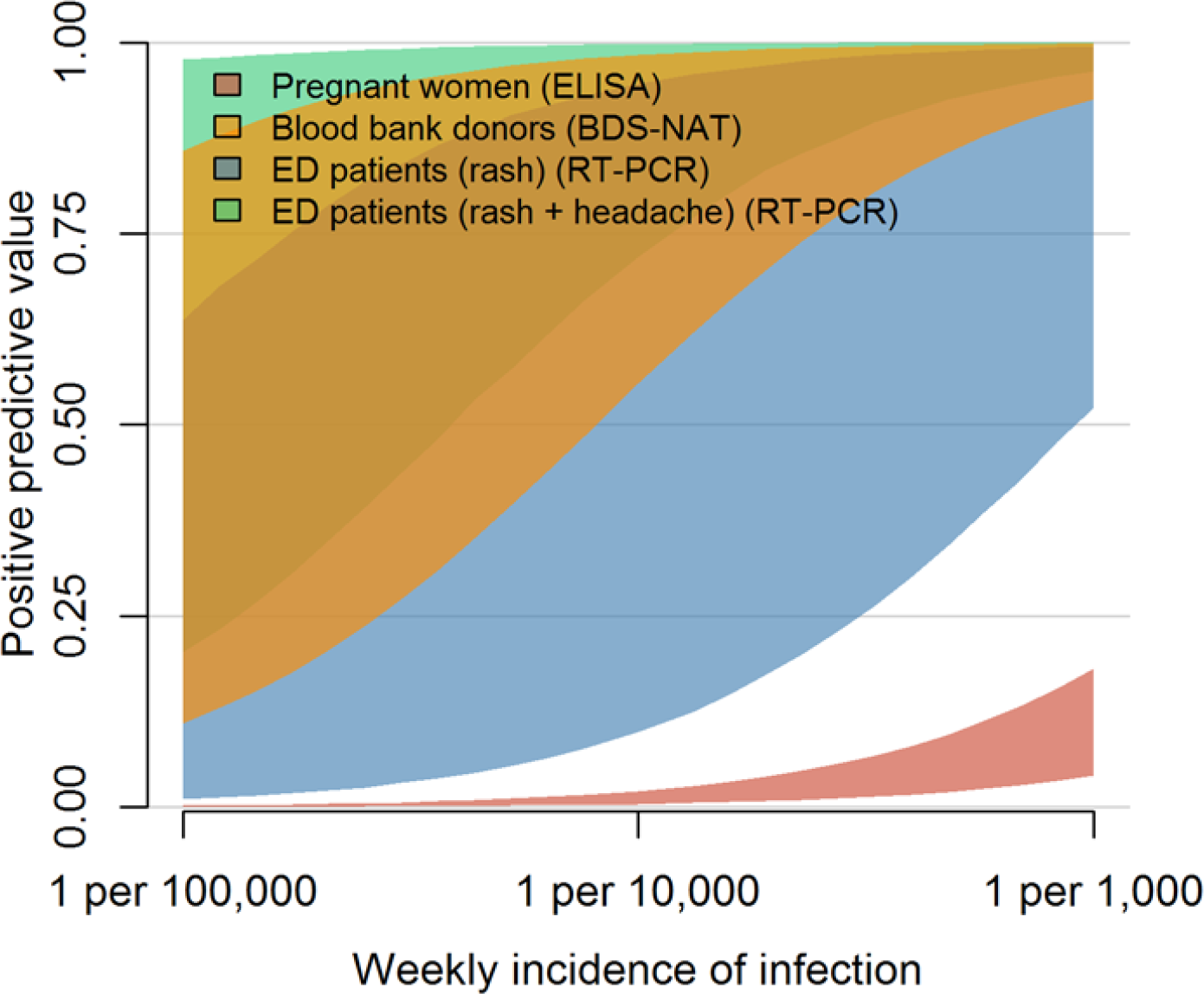
Positive predictive value for different surveillance strategies. This figure describes the positive predictive value (PPV) of a single positive test result under different surveillance strategies: testing asymptomatic pregnant women (red), blood donors (orange), ED patients exhibiting rash (blue) or ED patients exhibiting rash and headache (green). The overlapping bands represent 95% uncertainty intervals for the PPV of a positive test over a range of possible ZIKV incidences; Figure 3 shows 50% uncertainty intervals.

**Supplemental Table 1.** Estimates of syndrome prevalence

| **Parameter** | **Estimate among ZIKV patients seeking care** | **Estimate among ED patients** |
| --- | --- | --- |
| Fever | 51-53% | 3.7-6.7% |
| Headache | 65-67% | 2.7-3.6% |
| Rash | 86-87% | 4.3-6.1% |
| Arthralgia | 60-61% | 0.8-1.4% |
| Conjunctivitis | 16-17% | 0.1-0.3% |
| Arthralgia + Conjunctivitis | 12-13% | 0.0005-0.002% |
| Arthralgia + Fever | 35-37% | 0.001-0.003% |
| Arthralgia + Headache | 47-49% | 0.04-0.06% |
| Arthralgia + Rash | 53-54% | 0.1-0.3% |
| Conjunctivitis + Fever | 9.5-11% | 0.007-0.02% |
| Conjunctivitis + Headache | 12-13% | 0.2-0.4% |
| Conjunctivitis + Rash | 14-16% | 0.4-0.5% |
| Fever + Headache | 39-41% | 0.06-0.2% |
| Fever + Rash | 44-46% | 0.07-0.1% |
| Rash + Headache | 58-60% | 0.1-0.2% |
| Arthralgia + Conjunctivitis + Fever | 7.8-8.7% | 0-0.000001% |
| Arthralgia + Conjunctivitis + Headache | 10-11% | 0-0.001% |
| Arthralgia + Conjunctivitis + Rash | 11-12% | 0-0.001% |
| Arthralgia + Fever + Headache | 30-32% | 0.02-0.05% |
| Arthralgia + Fever + Rash | 31-32% | 0.0001-0.0004% |
| Arthralgia + Headache + Rash | 42-44% | 0.003-0.01% |
| Conjunctivitis + Fever + Headache | 7.9-8.8% | 0.002-0.005% |
| Conjunctivitis + Fever + Rash | 8.7-9.6% | 0.0007-0.002% |
| Conjunctivitis + Headache + Rash | 11-12% | 0.0004-0.001% |
| Fever + Headache + Rash | 35-36% | 0.007-0.02% |
| Arthralgia + Conjunctivitis + Fever + Headache | 6.9-7.7% | 0-0.000001% |
| Arthralgia + Fever + Headache + Rash | 27-28% | 0-0.000001% |
| Arthralgia + Conjunctivitis + Headache + Rash | 9.4-10% | 0-0.001% |
| Arthralgia + Conjunctivitis + Fever + Rash | 7.3-8.1% | 0-0.001% |
| Conjunctivitis + Fever + Headache + Rash | 7.4-8.2% | 0.001-0.004% |
| Arthralgia + Conjunctivitis + Fever + Headache + Rash | 6.4-7.3% | 0-0.000001% |
| Combination of two or more symptoms | 82-83% | 0.9-2% |
| Combination of three or more symptoms | 61-62% | 0.03-0.07% |
| Rash + one other symptom | 75-76% | 0.6-0.9% |

